# “Understanding Cortical Arousals during Sleep from Leg Movements: A Pilot Study.”

**DOI:** 10.1101/2021.09.14.460395

**Authors:** Kanika Bansal, Javier Garcia, Cody Feltch, Christopher Earley, Ryan Robucci, Nilanjan Banerjee, Justin Brooks

## Abstract

Leg movements during sleep occur in patients with sleep pathology and healthy individuals. Some (but not all) leg movements during sleep are related to cortical arousals which occur without conscious awareness of the patient but have a significant effect of sleep fragmentation. Detecting leg movements during sleep that are associated with cortical arousals can provide unique insight into the nature and quality of sleep in both health and disease. In this study, a novel leg movement monitor is used in conjunction with polysomnography to better understand the relationship between leg movement and electroencephalogram (EEG) defined cortical arousals. In an approach that we call neuro-extremity analysis, graph theoretic, directed connectivity metrics are used to interrogate the causal links between neural activity measured by EEG and leg movements measured by the sensors within the leg movement monitor. The leg movement monitor in this study utilizes novel capacitive displacement sensors, and a 9-axis inertial measurement unit to characterize leg and foot movements. First, the capacitive displacement measures more closely related to EEG-defined cortical arousals than inertial measurements. Second, the neuro-extremity analysis reveals a temporally evolving connectivity pattern that is consistent with a model of cortical arousals in which brainstem dysfunction leads to near-instantaneous leg movements and a delayed, filtered signal to the cortex. Neuro-extremity analysis reveals causal relationships between EEG and leg movement sensor time-series data that may aid researchers to better understand the pathophysiology of cortical arousals associated with leg movements during sleep.

## Introduction

Humans, unlike other animals, move almost exclusively upright on two legs that produce intricate patterns of leg movements from simple to complex, orchestrated mostly via the Central Nervous System (CNS). These processes are inhibited during restorative sleep, and dysfunction in this inhibitory neural circuitry, potentially caused by physiological irregularities, can manifest as specific leg movements while a person is asleep. For example, leg movements during sleep can reflect cortical and/or autonomic arousals, which represent significant changes in brain activity accompanied by, transitory increases in blood pressure and heart rate causing significant sleep disturbances [1–4].

Cortical arousals are theorized to occur as complex phenomena within the central nervous system that involves the cortex, thalamus, brainstem, and spinal cord. These arousals are currently observed and characterized by electroencephalography (EEG) that occurs as part of overnight polysomnography (PSG) or home sleep test (HST) studies. The sleeper is generally unaware of these arousals; however, these arousals can have a significant, negative impact [5–7].

Although the precise pathophysiology relating leg movements during sleep and cortical arousals remains unclear; since leg movements are associated with cortical arousals [8–10], their measurement is a critical, yet understudied measure of sleep and potential sleep disturbances. Current methods for measuring leg movements during sleep include actigraphy (accelerometer-only based devices) and EMG used in PSG; however, these devices give a limited view of the complex and sometimes subtle movements of lower extremities. For example, since actigraphy uses accelerometers, the leg must move enough to create a detectable signal above noise-levels and thus may miss more subtle, yet critical movements [11]. Furthermore, leg movements that do not involve the location of the accelerometers would be completely undetected. Alternatively, EMG can detect subtle movement (through the detection of voltage in muscle) but based on its location overlying the *anterior tibialis muscle*, can only identify leg movements associated with contraction of that muscle which controls dorsiflexion of the foot. Thus, any leg movements involving flexion at the knee or even some rotations of the foot would not be detected.

Given the hypothesized relationship between leg movements of sleep and brain activity, we developed a new analytic technique called neuro-extremity analysis that examines the effective connectivity between brain sensor (EEG) and leg movement sensors to better understand the temporal dependence of activity between the two. The goal of this approach was to better understand the mechanism behind leg movements associated with cortical arousals. Further, given the complexity of leg movements, we hypothesized that different types of leg sensors may show different strengths of leg movement/cortical connections.

In this pilot study, multiple devices were used to measure leg movements during sleep to evaluate their utility in detecting expert-labeled cortical arousal events and to evaluate their relationship with ongoing brain activity determined by EEG. The sensor data used in this study were the two current standards (EMG and accelerometry based), a textile capacitive based sensor and an IMU consisting of a 3-axis accelerometer, 3-axis magnetometers, and 3-axis gyroscope.

Our results show that in our small sample size of heterogeneous patients, the capacitive sensor data was most useful in prediction of cortical arousals and showed stronger associations with EEG oscillations than either of the more conventionally used sensor modalities. The temporal pattern of coordinated EEG and leg movement sensor activity changes observed in our results are consistent with the model of brainstem dysfunction causing leg movements during sleep that are associated with cortical arousals.

## Methods

### Participants and Data Collection

The experimental methods have been previously published [12]. All studies were approved by the accredited Institutional Review Board at Johns Hopkins University and all procedures followed the standards provided in the 1964 Declaration of Helsinki.

Briefly, subjects were introduced to the experimental protocol which included simultaneous PSG and leg sensor recordings while they slept. For this pilot study, PSG data from 2 adults (one RLS patient off medications and another normal subject with no sleep complaint) and 4 children (3 children with ADHD not on medication and one child with no sleep complaints) was collected simultaneously with a variety of other sensors on the legs.

### Materials

Sensors included 6 channels of EEG (O1, O2, C3, C4, F3, F4), 2 channels of EOG, and 2 channels of EMG activity. The bi-lateral *anterior tibialis* EMG and the video recording of legs and feet provided ground truth of leg movements during sleep. During these studies, participants wore the RestEaze device on both ankles. The RestEaze product is designed to gather high resolution leg movement information and includes the accelerometry, capacitance, gyroscopy,and magnetometer sensors. These recordings supported the algorithm development and are presented here.

### Cortical Arousal Markings

All PSG sleep measures used in the following analyses were obtained by visual scoring of the sleep studies by trained technicians with a careful review and adjustment of the results as needed by a board-certified sleep professional who was also an expert in leg movement and arousal recordings and measurements. All scoring followed the rules established by the American Academy of Sleep Medicine (AASM). Review of the EEG was conducted off-line after the data collection was completed. On average 80±25 (mean±SD) CAs were found for each participant. Importantly, these CAs were used in subsequent analyses and used as ground truth for CA.

### Analysis

EEG and leg sensor data were analyzed with two approaches to explore the relationship between leg movements and cortical arousals. The initial approach (Approach 1) was used to determine whether information gathered from a variety of sensors on the leg can predict a future cortical event. We use a traditional linear regression analysis and use a metric of explained variance to estimate successful prediction. Our next approach (Approach 2) was more exploratory and used traditional EEG connectivity techniques to investigate the statistical dependencies between the EEG time series and the leg sensor time series.

#### Approach 1: Linear regression

In our first analysis, we tested if the sensor data can be used to predict the occurrence of a cortical arousal using linear regression. In separate regression models, we used the coefficient of variation (CoV, standard deviation divided by the mean) of the data coming from different sensors for a fixed temporal window (t). These data included 3 of each capacitance, gyroscopy, accelerometer, and magnetometer sensors, 2 EMG sensors (Figure 1A), and an additional measure computed from capacitance sensors, which represented the sum of differential capacitance values across all three capacitance sensors (CDS). This quantity was included given the observed large fluctuation in the measured capacitance values.

**Fig. 1 -.**
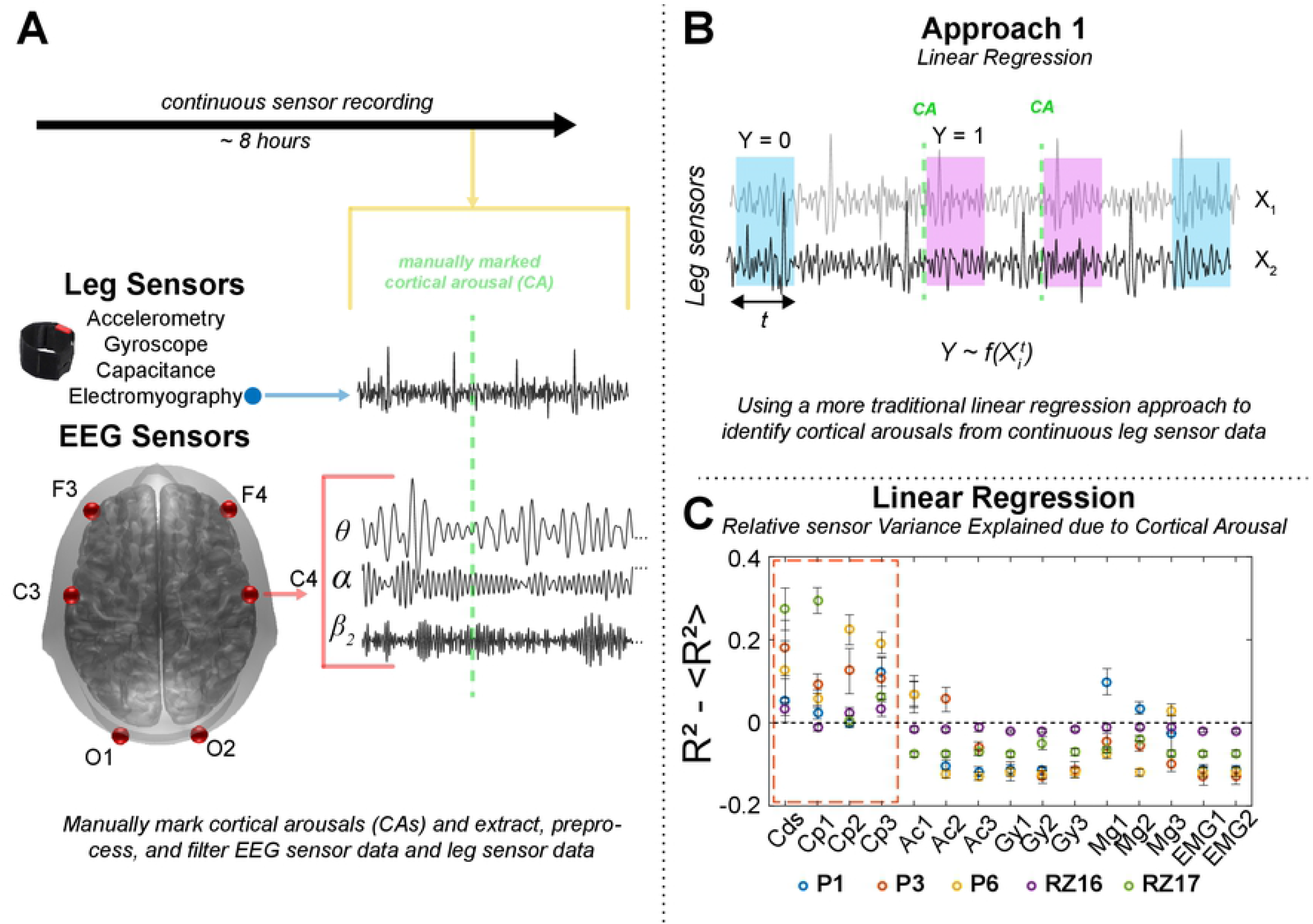
Linear Regression Analysis framework. A) Leg movement data during sleep was recorded from a variety of sensors including, accelerometer (Acc), gyroscope (Gyro), capacitance sensor (Cap), magnetometer (Mag), and from electromyography (EMG). In addition, brain activity was recorded using electroencephalography (EEG) from a couple of frontal (F), central (C), and occipital (O) regions. Arousals were marked manually during the processing of the data. For a part of the analysis, EEG data was filtered into different frequency bands. B) In our first analysis approach, we used regression models to test and compare if/how the data from different sensors can predict the occurrence of a cortical arousal. As a predictor, we used the coefficient of variation in the sensor data during a temporal window (t) following an arousal (response variable Y = 1) and no arousal (Y = 0). C) To compare across sensors, here we show the standardized R^2^ for each sensor and subject, which was calculated by subtracting the mean R^2^ values across all the sensors for a given subject. Error bars represent the standard deviation across 20 regression trials where the equal number of no-arousal windows were randomly chosen (see Methods). We observed that use of capacitance sensors was better than use of other sensors in differentiating arousal events from no arousal events.

Shown as Approach 1 in Figure 1B, regression was done on a binary dependent variable (*Y*) which identified if the CoV was calculated for the data *t* seconds after a CA (*Y* = 1) or for a randomly selected window of the same length which did not follow the occurrence of a CA (null data, *Y* = 0). For every CA observed, a null window was identified to be included in the model. For the robustness of this analysis, we conducted 20 regression trials for every stream of the sensor data, and in each trial, required null data windows were selected randomly from the entire duration of the recording.

Regression analysis was done separately for each individual and to compare different sensors as predictors, we used F-statistics, p-values, and R^2^. Using initial testing, we found a window (*t*) of 15-30 seconds to be optimum for the regression models. As a representative comparative metric, we used a standardized R^2^ (Figure 1C) for each sensor and subject, which was calculated by subtracting the mean R^2^ values across all the sensors for a given subject.

#### Approach 2: Neuro-extremity Analysis, Granger Causal Modelling

Our neuro-extremity analysis method calculated time-varying asymmetries between fluctuations in oscillatory activity within the brain and the fluctuations in a variety of sensors placed on the lower extremity. We employed a Granger causal framework to investigate the predictive relationship between the brain activity in the temporal proximity of a clinically defined cortical arousal and sensors specially designed to capture leg movements from a variety of high-resolution sensors. Previous research established that theta (5-8Hz) rhythms are uniquely tied to sleep and locomotion across species and are used in the scoring of cortical arousals according to v2.6 of the AASM scoring manual. Therefore, in our second approach we focused on theta rhythms.

Granger causal analyses have been used in EEG analyses for some time and have proven a fruitful endeavor to understand how coordinated neural activity may give rise to cognition.^13,14^ We have recently expanded these analytical techniques not only to understand connectivity within the brain, but also to understand how fluctuations in neural activity might explain other variables, like continuous motor behavior.^15^ Largely following this framework, we have expanded these methods to understand: (1) which features of leg movements and cortical arousals, as measured by sensors placed on the leg and scalp, are closely intertwined, (2) the temporal limitations and extent of this relationship, and (3) the extent to which these neuro-extremity relationships may be further explored.

This approach is mainly completed within a few steps that include: (1) preprocessing the EEG and leg sensor data, (2) determining the model parameters, (3) fitting the model, (4) estimating partial directed coherence, (5) statistical testing, and (6) evaluating asymmetry between the brain-predicting-leg and leg-predicting-brain estimations. These steps are detailed below and are roughly depicted in Figure 2 (A,B,C,E).

1. Because the methods require continuous time series and a generally large amount of time segments to build a predictive model of neuro-extremity connectivity, special care must be considered to discard any perturbations in the signal that might be driven by nuisance factors (e.g., EEG artifact like jaw clenches, poor sensor contact, etc). Due to some body movement over the > 8hr recording and the limited dataset within this pilot study, we used an artifact subspace removal (ASR) on the electrodes, which decomposes the EEG into principal components in overlapping sliding windows to identify and reject high amplitude components. This method has previously shown to be highly successful in naturalistic EEG data collections.^16^ After this continuous correction of the time series, EEG data were filtered with a low-order (3) butterworth zero-phase filtering scheme. Leg sensor data was not filtered or processed so as to capture any and all possible features from the sensors that might be useful in prediction of (or predicted by) EEG activity surrounding a cortical arousal. Finally, the EEG data were epoched into a 7 second window surrounding the cortical arousal. The leg sensor data was epoched into 7 second windows through the duration of 6 mins surrounding the cortical arousal to understand the temporal dynamics of the prediction. Separate AR models were estimated for each of the leg sensor epochs.
2. Next, the ARfit toolbox^17^ was used to implement a stepwise least squares estimation of multivariate auto-regressive (AR) model at a variety of model orders (5-30), and the Schwarz Bayes information criterion (SBC) across the model orders was inspected to find the optimal model order (20) for these complex neuro-extremity relationships. The relatively high model order was used to capture as many possible temporally dependencies as might be available in the time series.
3. Finally, we calculated the partial directed coherence (PDC), a measure that describes the direction of information flow based on a multivariate decomposition of the partial coherences within complex time varying signals.^18,19^ This resulted in a 21 x 21 matrix for all pair-wise EEG and leg sensor directional relationships.
4. The PDC estimates, on their own, imply the information flow between two time series derived from a metric that signifies how a time series may predict a future fluctuation in another time series, but they could also signify some inherent predictable feature in both signals. To determine the robustness of the PDC, a null distribution of PDC values was created by randomly rotating the time series of the EEG and leg data. PDC was estimated in the shuffled data, and then all estimates were aggregated. Only the PDC values above (or below) the 95% confidence interval of the null distribution mean were considered. Figure 2C steps through this process showing the mean PDC across all sensors (left panel), then a count of time points that are significantly above (or below) the 95% confidence interval of the null distribution (middle panel), and then finally, the mean PDC when only considering the significant edges (right panel).
5. Lastly, when estimating the directed connectivity in this dataset, we studied the time course of PDC, specifically to understand the asymmetry in the prediction. We estimated the *asymmetry index (AI)* as the difference normalized by the sum of each pairwise sensor relationship. If the AI is < 0, then it suggests more that leg sensor measurements can be used to predict the EEG measurements than the EEG sensor measurements can be used to predict the leg sensor measurements. If the AI is > 0, then the opposite is true. After the AI was estimated across the time course, the time-evolving asymmetry matrix was subjected to the same statistical thresholding procedure in (4) and only those significant PDC values were used to estimate the average asymmetry 1 minute before the CA and 1 minute after the CA, now known as CA_pre_ and CA_post_.

**Figure 2 -.**
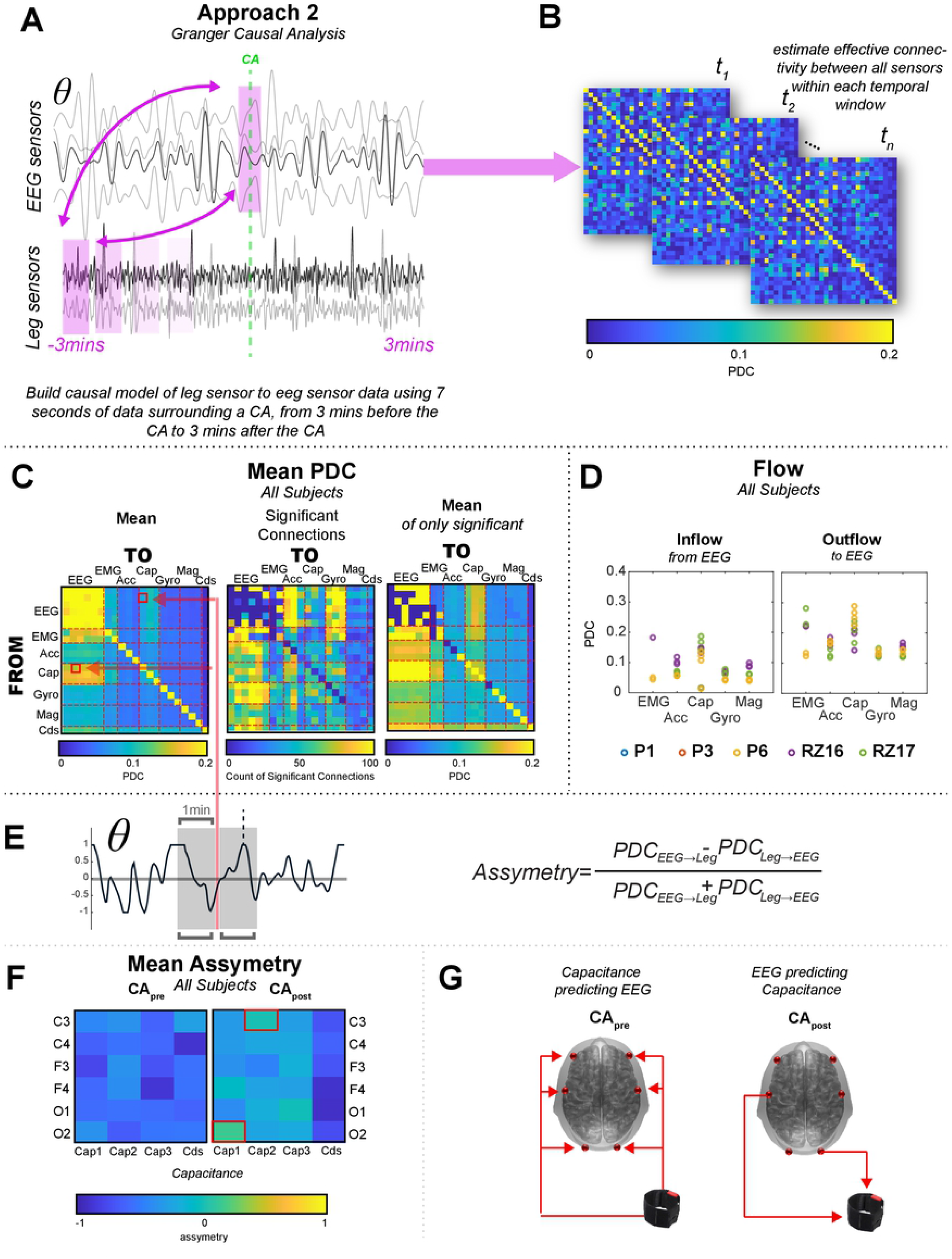
Approach 2: Neuro-extremity Analysis, Granger Causal Approach: A) Visual depiction of the approach, where (after substantial preprocessing) causal models of 7 secs of sensor data were built in several windows from 3 mins before the CA to 3 mins after. Importantly, only the leg sensor data was changed in these windows, the CA EEG data was never shifted in time. B) For each time window, partial directed coherence (PDC) was estimated between each pairwise sensor and an asymmetrical matrix was produced that represented the directional relationships inferred via PDC. C) Summary PDC matrices across all subjects. Mean PDC across all participants, without statistical thresholding (left panel), number of significant time points across all participants and time windows for each cell of the PDC matrix (middle panel), and the mean PDC across all participants when only considering the significant PDC cells (right panel). D) Inflow and Outflow of the leg sensor types within the study, calculated by the sum of significant EEG PDC values. E) Asymmetry Index (AI) was calculated to determine the bias in directionality in the PDC values within the 2 mins surrounding the CAs. A sample time course of the AI is shown for the EEG-theta band. F). While the CA_pre_ is dominated by the relationship of the Capacitance sensors predicting the CA, the CApost interval is less uniform, with certain Capacitance/EEG sensor pairings showing the opposite. These pairs are highlighted with red. G) Visual display of the Cap-to-EEG relationship before the CA (CA_pre_) and the relationship of the two notable exceptions that show higher PDC from the EEG sensors (C3,O2) to the Cap sensors (right panel) after the CA (CA_post_).

## Results

### Linear Regression suggests capacitance sensor can identify CAs

We used linear regression models for each sensor to test if the coefficient of variation in the sensor time series can dissociate arousal events from non-arousal events (Figure 1B, also see Methods). We assessed model F-statistics, p-values, and the R^2^ values to test how each sensor performed. For each sensor stream the mean and standard deviation of the R^2^ across all channels was displacement capacitative: 0.2 ± 0.12; accelerometers: 0.06 ± 0.072; gyroscope 0.0084 ± 0.008; magnetometers 0.065 ± 0.07; and EMG 0.005 ± 0.0026. In Figure 1C, we show the standardized R^2^ calculated for all the sensors across different subjects to summarize our findings. We calculated this quantity by subtracting the mean R^2^ value observed across all the sensors from the R^2^ value observed for the given sensor for each subject and regression trial. We found that the capacitance sensors performed the best and separated the arousals from non-arousal reasonably well.

### Neuro-extremity analysis reveals special role of capacitance in EEG signal prediction

As a more exploratory approach, to support the analysis in Approach 1, we implemented a neuro-extremity analysis technique that employs an EEG connectivity approach, partial directed coherence (PDC) to inspect the directional information flow between all pairwise sensors. Figure 2 visually describes this analysis technique, which is largely derived from a previous neuro-behavioral analysis used to describe the relationships between the brain and continuous behavioral changes.15 This resulted in a series of PDC matrices that represented how well one signal may be used to predict another, from 3 minutes before the cortical arousal to 3 minutes after the cortical arousal. To determine whether the value of each cell of the PDC matrix was significant, we estimated a null distribution and only considered values that were higher than the 95% confidence interval surrounding the mean of the null distribution, and then only considered these values in the overall mean (Figure 2C, right panel). As is observed in Figure 2C, a structured pattern emerges for each of the panels. First, the overall mean PDC values, without statistical thresholding (Figure 2C, left panel), shows high mean PDC for connectivity between the EEG sensors. Capacitance (Cap) and EMG sensors also show relatively high mean PDC with directional information flow from the leg sensors to the EEG sensors, without much spatial specificity. Next, the number of significant connections across all 5 subjects and time windows (Figure 2C, middle panel: 6 total minutes, 241 windows), suggests many of the cells representing the information flow between the EEG sensors and leg sensors show many significant time windows in the cells of the time varying PDC matrix, as compared to the lack of significant pairs within the EEG sensors alone. Finally, when only considering the significant windows, the mean PDC from the Capacitance sensors to the EEG sensors appears to be higher than the alternative direction, suggesting the temporal fluctuations in the leg sensors are more predictive of the EEG signal fluctuations than the alternative.

Finally, to confirm the specificity of the results shown in the mean PDC matrix (Figure 2C, right), we next summed the significant PDC values over all 6 EEG sensors PDC and inspected the inflow and outflow of each type of leg sensor for each subject. In other words, we were interested in how well each leg sensor measurement was influenced or predicted using the EEG sensors (i.e., inflow) and vice versa (i.e., outflow). Supporting the findings in Figure 2C, the Cap sensors display the highest overall inflow and outflow with a couple of exceptions in which a subset of the EMG sensors in 2 participants also displayed similarly high inflow and outflow. Interestingly, 2 of the 3 pediatric patients did not have time windows that survived the null model comparison, as the variance in the statistical dependencies between all sensors in the null model was higher than that in the other subjects.

### Asymmetry in neuro-extremity analysis suggests a new pathway to investigate brain/body relationships

While the previous summary of results referred to the specificity in EEG-leg sensor prediction, our final step in the neuro-extremity analysis, directly compares the EEG-leg sensor PDC values across time. To perhaps shed light onto the observed relationships and taking advantage of the temporal information present in the time-evolving PDC matrices, we next derived an asymmetry index (AI), which directly estimates the directional bias in PDC values. For illustrative purposes, we only inspect the most robust Cap sensor and specifically ask whether there are pairwise relationships that may change their directional bias in windows before (CApre) and/or after (CApost) the CA. Figure 2F is a visualization of the average asymmetry (considering only significant time windows), of the individual EEG sensors to all of Cap sensors, including the differential sum of all 3 Cap sensors (Cds). We observe that overall, there is a substantial bias in the leg sensors predicting the temporal dynamics of the EEG sensors, and this relationship is especially apparent in the 1 min before a CA (Figure 2F, compare left and right panels); however, there are a couple of noticeable exceptions, in central and posterior electrodes (C3, O2) for a subset of the Cap sensors. Figure 2G pictorially represents these relationships, with red arrows representing the direction of prediction. These results add to the previous findings that showed perhaps special features in the dynamics of Cap sensors as they can predict CAs and provide a framework to further study brain-corporeal investigations.

## Discussion

In this pilot study textile capacitive sensor measurements of the lower extremity were more valuable in discriminating cortical arousals than other sensor modalities and further, these data were more closely related to EEG-theta oscillations near the cortical arousal events. Collectively, these results (1) provide preliminary support that complex leg movements during sleep are associated with cortical arousals and (2) provide preliminary evidence of causal connections (neuro-extremity analysis) between these leg movements and the brain during cortical arousals.

Leg movements during sleep can be significant biomarkers for cortical arousals that reflect significant fragmentation in homeostatic sleep mechanisms and are implicated in multiple sleep disorders,^20^ attention deficit hyperactivity disorder,^21^ neurodegenerative disease,^22^ and cardiovascular disease.^23^ Despite these significant relationships, it is still unknown whether this reflects (i) a process in which dysfunction in the brainstem leads to an immediate movement of the legs with delayed processing through the thalamus to cortex or (ii) somatosensory feedback of the leg movement to cortex, or perhaps a combination of the two mechanisms. Indeed, the emergence of a leg movement during sleep is a complex process involving numerous muscles of the lower extremity and several cortical and subcortical structures.^24^ Because of this complexity, there is significant variability in the type and duration of leg movements during sleep. In this study, using capacitive sensors, we were able to capture and model this complexity in relationship to cortical arousals and EEG dynamics related to those arousals. The first analysis revealed that the relative variability accounted for in detecting cortical arousals was highest with the capacitive sensors (Figure 1C). Similarly, the neuro-extremity analysis that examines the connectivity between EEG-theta activity and leg movement sensor data, revealed stronger effective connectivity between the capacitive sensors and EEG-theta oscillations than other sensor modalities (Figure 2C and 2D). Capacitive sensors can capture more diverse leg movements during sleep than the other sensors. By design, the capacitive sensors capture relative changes in the electromagnetic field that are generated through changes in lower extremity position. This enables the sensor to capture numerous types of leg movements including those that do not involve the anterior tibialis (muscle used during PSG/EMG studies) nor the physical movement of the ankle itself (e.g. dorsiflexion of the great toe) used by actigraphy (accelerometry based devices). Thus, the present results that show a stronger relationship between these data and EEG-theta activity may be able to capture a more general movement circuit between leg movements and the brain. The other leg sensors used in this study detect leg movements during sleep but are more specific to particular types of movements. For example, the EMG channels used here (and in common practice for PSG) will only detect activation of the anterior tibialis muscle which controls dorsiflexion of the foot but may miss the other movements (e.g. flexion at the knee). Similarly, actigraphy requires that the sensors physically move sufficiently above baseline noise levels for leg movements to be detected. Thus, subtle movements measured by actigraphy may not show relationships to cortical arousals and neural activity that are detected by capacitive sensing.

The neuro-extremity axis analysis reveals an interesting pattern of effective connectivity. The temporal relationship between leg movements and EEG-theta changes shows significant effective connectivity from the leg movement to the cortex before the manually demarcated cortical arousal (Figure 2F). This implies that leg movements precede detectable changes in EEG before the cortical arousal. After the arousal, there is, similarly, significant outflow from the leg-sensors to EEG, but several EEG-leg sensor pairs that also show the opposite directionality (effective connectivity from EEG to the leg movement time series) emerge during this time.

Collectively, we interpret these findings as representative of a theory suggesting that certain leg movements during sleep (those associated with cortical arousals), originate in the brainstem and signal in two pathways with two different delays.^25^ One pathway, from the brainstem through the spine and to the muscles of the leg which is relatively quick to generate a leg movement. It originates in the brainstem and involves subcortical structures including central pattern generators (CPGs) that are otherwise inhibited during sleep.^26^ The other pathway, from the brainstem through the thalamus and to the cortex is slower relative to the brainstem to leg movement pathway and results in a cortical arousal following a leg movement. The brainstem to cortex pathway is involved with sleep/wake balance and this is likely involved in the sleep fragmentation, cortical arousal associated with the particular leg movements studied in this manuscript.^27,28^

The current results are consistent with this model. From a temporal perspective, the inciting event is thought to originate in the brainstem in which CPGs create prototypical leg movements during sleep that begin very early in the process. Simultaneously, a delayed signal from the brainstem through the thalamus and to cortex is transmitted resulting in detectable changes in EEG. This signal is detected as an effective connection between the leg movement and EEG changes in Figure 2F (left) which may be related to the cortical arousal event. As the legs continue to move, the associated cortical activity changes (as measured by EEG) resulting in effective connectivity from EEG to leg movements as shown in Figure 2F (right). This conceptual model is captured in Figure 3.

**Figure 3 -.**
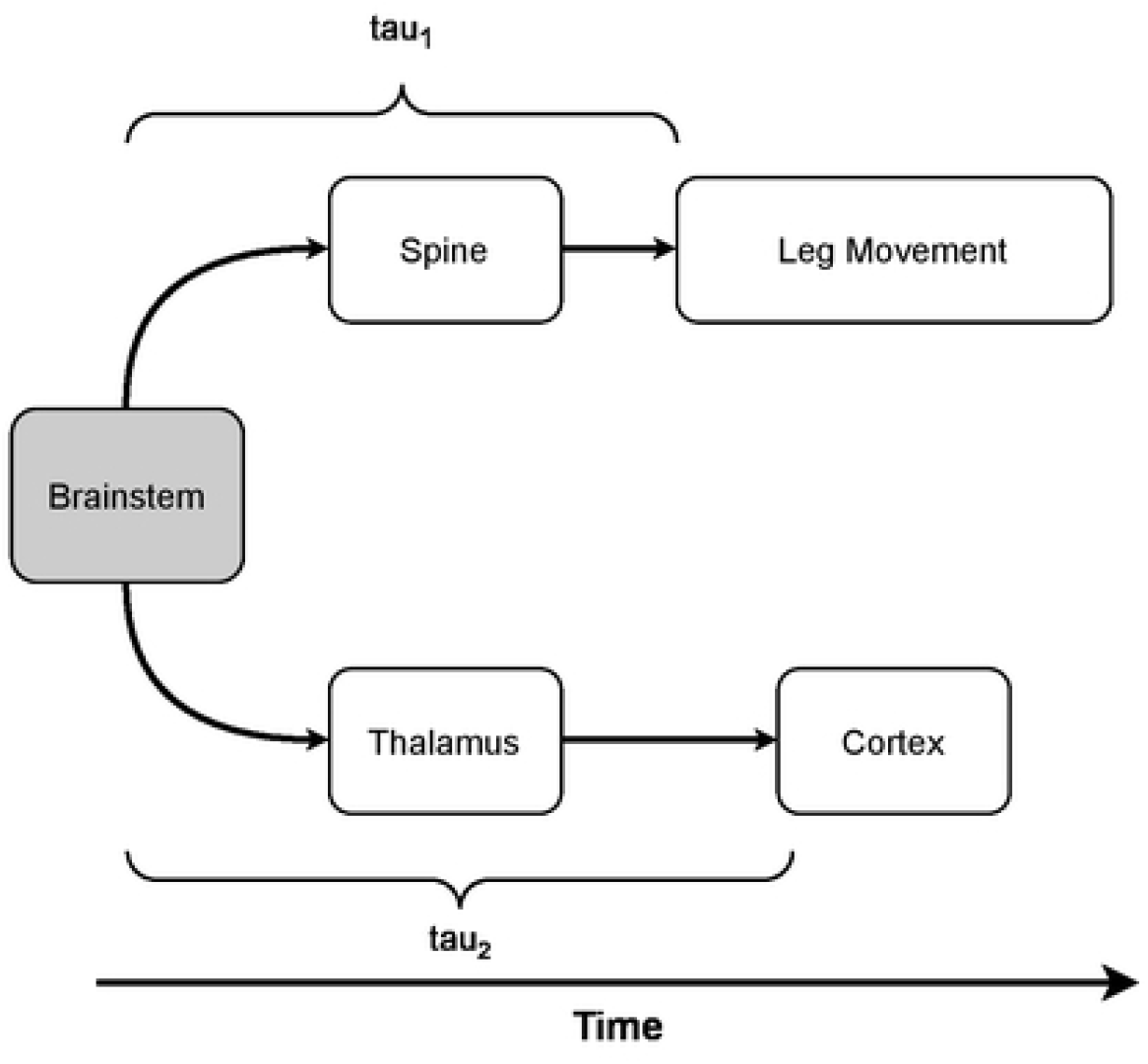
A conceptual model of LMS associated with arousals: to show the timing between CNS areas involved in the leg movements during sleep associated with cortical arousals. Tau_1_ represents the delay from the brainstem event to the beginning of the leg movement which is quick. Tau_2_ represents the delay between the brainstem event and its transmission to the cortex which is slow.

Other studies have shown different temporal patterns of EEG and leg movement activity. Specifically, in periodic leg movement disorder a study demonstrated that delta rhythms precede PSG (EMG-derived) estimates of leg movements which the authors conclude as representative of a cortical origin of periodic leg movements during sleep in Periodic Leg Movement Disorder (PLMD) patients. ^29^ Another study has demonstrated similar findings to ours in Restless Leg Syndrome (RLS) patients where significant alpha and beta spectral power changes were observed after the initiation of leg movements with no change in delta range. ^30^ There are a few reasons for this discrepancy. First, these two studies used patients with different diagnoses (PLMD vs RLS) which may have different underlying pathophysiology and are associated with different sleep characteristics.^31^ Second, our approach to the measurement of leg movement is unique. Instead of using EMG of the *anterior tibialis* we used a textile based capacitive sensor. This sensor will detect more leg movements than an EMG probe and thus is better suited to capture more complex leg movements that involve different muscles.

Preliminary evidence collected by our group suggests that leg movements begin earlier and last longer than those detected by EMG alone (Brooks, Feltch et al, under review).

## Conclusions

Leg movements during sleep associated with cortical arousals are complex and poorly understood phenomena. Our preliminary research has provided some additional insight by demonstrating that (1) capacitive sensor data can provide an improved ability to detect cortical arousals based on leg movement data and (2) proposes a neuo-extremity analysis to estimate the temporal dynamics in the EEG-leg movement relationship. Initial implementation of this approach shows consistency with models of brainstem dysfunction as a potential source of leg movements during sleep associated with cortical arousals.

There are a few limitations to consider with this study. First, the sample size is quite small and heterogenous and additional data is needed to further clarify the physiology of leg movements during sleep. Second the sparse locations of the EEG sensors used in this study is poor, thus prohibiting clear interpretation of the functional areas of cortex involved. Finally, additional characterization is needed to understand the precise muscle activations (and subsequent movements) associated with the capacitive time series data. Future research is needed to clarify these points.

## Acknowledgements

We are exceptionally grateful for the insights and ideas provided by the late Richard Allen, Ph.D. We thoroughly enjoyed reviewing earlier versions of this manuscript with him and are excited to continue our understanding of sleep pathology based on his work. We are also grateful to our patients for their time.

## Data Availability

The data underlying this article will be shared on reasonable request to the corresponding author.

